# Dysfunction of transfer RNA modifications in inflammatory bowel disease

**DOI:** 10.1101/2022.05.18.492467

**Authors:** Jilei Zhang, Yongguo Zhang, Yinglin Xia, Jun Sun

## Abstract

**Backgrounds and aims:** Transfer RNA (tRNA) is the most extensively modified RNA in cells. Queuosine (Q)-modification is a fundamental process for fidelity and efficiency of translation from RNA to proteins. In eukaryotes, tRNA-Q-modification relies on the intestinal microbial product queuine. However, the roles and potential mechanisms of Q-tRNA modifications in IBD are unknown.

**Methods:** We explored the Q-tRNA modifications and expression of Q tRNA ribosyltransferase catalytic subunit 1 (QTRT1) in patients with IBD by investigating human biopsies and reanalyzing datasets. We used colitis models, organoids, and cultured cells for loss- and gain-of-function studies to investigate the molecular mechanisms of Q-tRNA modifications in intestinal inflammation.

**Results:** QTRT1 expression was significantly downregulated in ulcerative colitis and Crohn’s disease patients. The four Q-tRNA-related tRNA synthetases (asparaginyl-aspartyl-, histidyl-, and tyrosyl-tRNA synthetase) were decreased in IBD patients. This reduction was further confirmed in DSS-induced colitis and IL10-deficient mice. Reduced QTRT1 was significantly correlated with cell proliferation and intestinal junctions, including downregulated β-catenin and Claudin-5 and upregulated Claudin-2. These alterations were confirmed in vitro by deleting QTRT1 from cells. Queuine treatment significantly enhanced cell proliferation and junction functions in cell lines and human colonoids. Queuine treatment also reduced inflammation in epithelial cells. Moreover, altered QTRT1-related metabolites were found in human IBD.

**Conclusion:** tRNA modifications play an unexplored novel role in the pathogenesis of intestinal inflammation by altering epithelial proliferation and junctions. Investigations on tRNA modification will uncover novel molecular mechanisms for potential prevention and therapy for IBD.

## Introduction

Eukaryotes acquire queuine as a nutrient factor from intestinal microbiota or from diet. In eubacterial and eukaryotic cells, queuine (q) is found as the sugar nucleotide queuosine (Q) at the wobble anticodon position of tRNA for the amino acids tyrosine, histidine, asparagine, and aspartic acid ^1^. Queuine produced by microbiota is taken up from the colonic lumen into enterocytes through cellular uptake mechanisms followed by incorporation into the wobble anticodon position of the four tRNAs by a heterodimeric enzyme encoded in the genome ^2-4^. This enzyme is a tRNA-guanine transglycosylase complex composed of the catalytic subunit QTRT1 and the noncatalytic subunit queuine-tRNA ribosyltransferase domain containing 1 ^5^. The physiological requirements for queuine and queuine-modified tRNAs have been documented in broad subjects for over four decades, establishing their relationships to development, proliferation, differentiation, metabolism, cancer and tyrosine biosynthesis in eukaryotes and invasion in pathogenic bacteria ^2^. Starvation for queuine and/or Q-deficiency in tRNA can cause changes in the pattern of protein synthesis. In addition, queuine deficiency may interfere with other nutrient factors, including vitamin B12, tetrahydrobiopterin, and epidermal growth factor ^1, 6^. Growing evidence indicates that tRNA modifications play important roles in human diseases, e.g., type 2 diabetes ^3^. Q depletion led to endoplasmic reticulum stress in mouse liver ^7, 8^. Q-tRNA modifications are dynamic and highly variable depending on the developmental stages and species type ^1, 7^ and tumorigenesis ^9^. However, the health consequences of disturbed availability of queuine and altered Q-tRNA modification remain to be investigated. The effects and mechanisms of Q-tRNA in digestive diseases are unknown.

IBD is a chronic disease that primarily affects the intestine, including Crohn’s disease (CD) and ulcerative colitis (UC). The etiology of IBD has been characterized as chronic intestinal inflammation resulting from many factors, including micronutrients and epigenetic mechanisms via DNA methylation and noncoding RNA ^10, 11^. Tissue barriers play an essential role in intestinal health ^12^. An increase in the tight junction protein Claudin-2 and a decrease in Claudin-5 may lead to the conversion of tight into leaky tight junctions in IBD patients ^13-17^. Our recent study indicated that Q-tRNA modification not only depends on the gut microbiome but also affects tight junctions, which regulate intestinal permeability in the development of breast cancer ^14^. However, the mechanisms by which Q-tRNA modifications are associated with intestinal functions in human IBD have never been investigated.

In the current study, we focus on addressing fundamental questions in IBD: is Q-tRNA modification involved in human IBD, and how does dysfunction of Q-tRNA modification contribute to chronic inflammation? We found that QTRT1 expression was significantly downregulated in UC and CD patients. Four Q-tRNA-related tRNA synthetases (e.g., asparaginyl, aspartyl, histidyl, and tyrosyl tRNA synthetase) were also decreased in IBD patients. Decreased QTRT1 expression was confirmed in experimental DSS-induced colitis and IL10-deficient mice. We investigated the impact of QTRT1 on tight junctions and proliferation in human organoids, cell lines, and colitis models. Insights into Q-tRNA modification in regulating barrier functions would facilitate the development of targeted interventions for IBD through tRNA modifications.

## Materials and Methods

(Detailed in the supplementary document)

### Human IBD Datasets reanalysis

We took advantage of microarray data reported in the Gene Expression Omnibus (GEO) repository (https://www.ncbi.nlm.nih.gov/geo/). For metabolic compound analyses, we revisited the datasets available in Metabolomics Workbench (https://www.metabolomicsworkbench.org/).

### Animals and animal models

Wild-type C57BL/6 and IL10 knockout (IL 10^−/-^) mice were purchased from Jackson Laboratory (Bar Harbor, Maine, USA). The mouse colitis model was induced by administering DSS, as previously described ^18^. Experiments were performed on 8- to 12-week-old mice (both male and female). Animals were provided the same and consistent conditions and utilized in accordance with the UIC Animal Care Committee, the Office of Animal Care and Institutional Biosafety (OACIB) guidelines, and the animal protocol (number ACC 18-179).

## Results

### Downregulation of QTRT1 in human IBD patients

To study the role of tRNA-Q modification in IBD patients, we investigated the expression of QTRT1 in IBD patients, including both UC and CD patients, and healthy controls by revisiting datasets in the NCBI GEO database (https://www.ncbi.nlm.nih.gov/geo). In the dataset with accession number GSE9452 ^19^, biopsies from the descending colon were collected from 14 UC patients with macroscopic signs of inflammation and 5 patients without any inflammation signs. We found that the mRNA expression of QTRT1, the subunit of the key enzyme for queuosine-containing tRNA modification, was significantly downregulated in the intestinal mucosa of the patients with UC symptoms compared with the non-UC patients (p<0.05) (Figure 1A). Similarly, when we reanalyzed the GSE83448 dataset ^20^, which has ileal biopsy samples from 39 patients with CD symptoms and 14 patients without any intestinal symptoms, QTRT1 expression was also found to be significantly downregulated in CD patients compared with controls (p<0.01) (Figure 1B). Since these downregulations were found at the mRNA level, we further investigated the alterations in QTRT1 protein expression in the biopsy colon tissue from IBD patients by immunohistochemistry (IHC) staining. In the UC patients, the QTRT1 staining intensity in the intestinal biopsy tissue was significantly reduced compared with controls (p<0.01) (Figure 1C). Meanwhile, the expression of QTRT1 was also found to be significantly downregulated in the intestinal tissue of CD patients compared with normal controls (p<0.01). However, no staining was observed in our IgG negative control for IHC staining (Figure 1D). More importantly, our findings of QTRT1 alteration at the protein level were consistent with the mRNA level in IBD patients. Taken together, we found that QTRT1 was downregulated at the mRNA and protein levels in the intestinal mucosa of IBD patients, including UC and CD patients.

**Figure 1.**
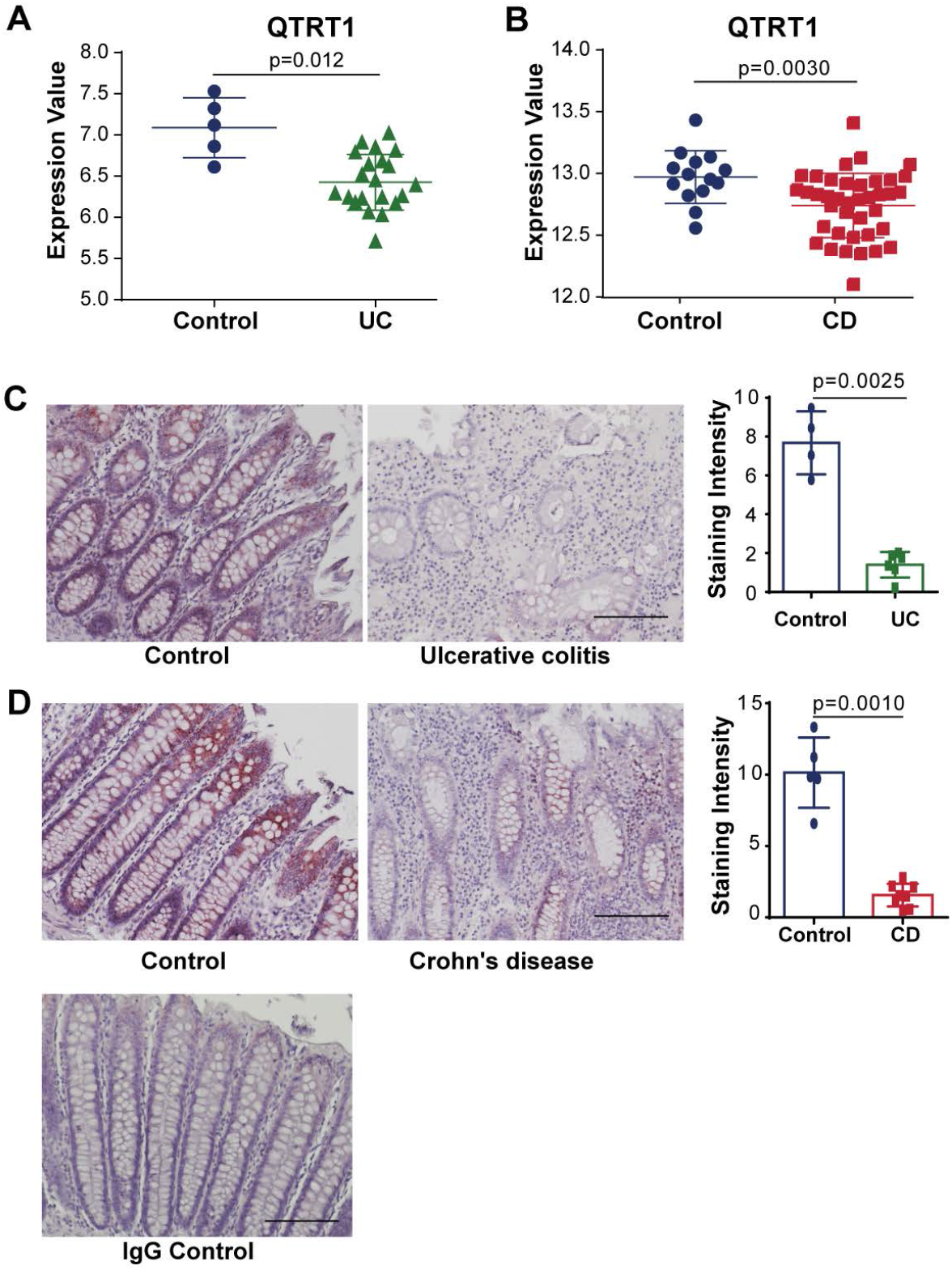
Expression of QTRT1 in human Crohn’s disease and ulcerative colitis patients. (**A**) Graphs of RNA-seq gene expression levels of QTRT1 in human colon biopsies of UC (n=14) patients and controls (n=5) from published databases. The data of UC patients were from NCBI GEO datasets (https://www.ncbi.nlm.nih.gov/geo) with accession number GSE9452. Data are shown as the mean ± SD, Welch’s t test. (**B**) Graphs of RNA-seq gene expression levels of QTRT1 in human colon biopsies of CD (n=39) patients and controls (n=14) from published databases. The data of CD patients were from NCBI GEO datasets (https://www.ncbi.nlm.nih.gov/geo) with accession number GSE83448. Data are shown as the mean ± SD, Welch’s t test. (**C**) Representative IHC staining of QTRT1 in colon tissue from ulcerative colitis (n=6) patients and controls (n=4). The scale bar is 300 μm. Semiquantitative analysis was performed on immunohistochemistry staining using ImageJ Fiji. Data are shown as the mean ± SD, Welch’s t test. (**D**) Representative IHC staining of QTRT1 in colon tissue from CD patients (n=8) and controls (n=5). The scale bar is 300 μm. Semiquantitative analysis was performed on immunohistochemistry staining using ImageJ Fiji. Data are shown as the mean ± SD, Welch’s t test. IgG was used as a negative control for all IHC staining.

### Reduction of QTRT1 in colitis mouse models

We then further investigated the expression of QTRT1 in colonic cells from IBD mouse models, including a DSS-induced colitis mouse model and an IL-10-deficient mouse model (Figure 2). We first evaluated the mRNA expression of QTRT1 in colon tissue using real-time PCR. We found that the mRNA expression of QTRT1 was significantly downregulated in the colon tissue from the DSS-treated mice compared with the nontreated control mice (p<0.01) (Figure 2A). Then, we evaluated the protein expression of QTRT1 in the colon tissue of DSS-induced colitis mice via western blot analysis. Clearly, the expression of QTRT1 in the DSS-treated mice was significantly suppressed compared with that in the controls (p<0.001) (Figure 2B). To further verify the protein expression of QTRT1 in the colitis mouse model, immunofluorescence staining was performed on colon tissue collected from DSS-treated mice. Inflammation and intestinal damage were found in the colon tissue from the DSS-treated mice (Figure 2C). Additionally, the expression of QTRT1 (fluorescence intensity) in the mice with colitis was significantly suppressed compared with that in nontreated control mice (p<0.01) (Figure 2C). However, there was no fluorescence staining in the IgG negative control used in the whole process of IF staining.

**Figure 2.**
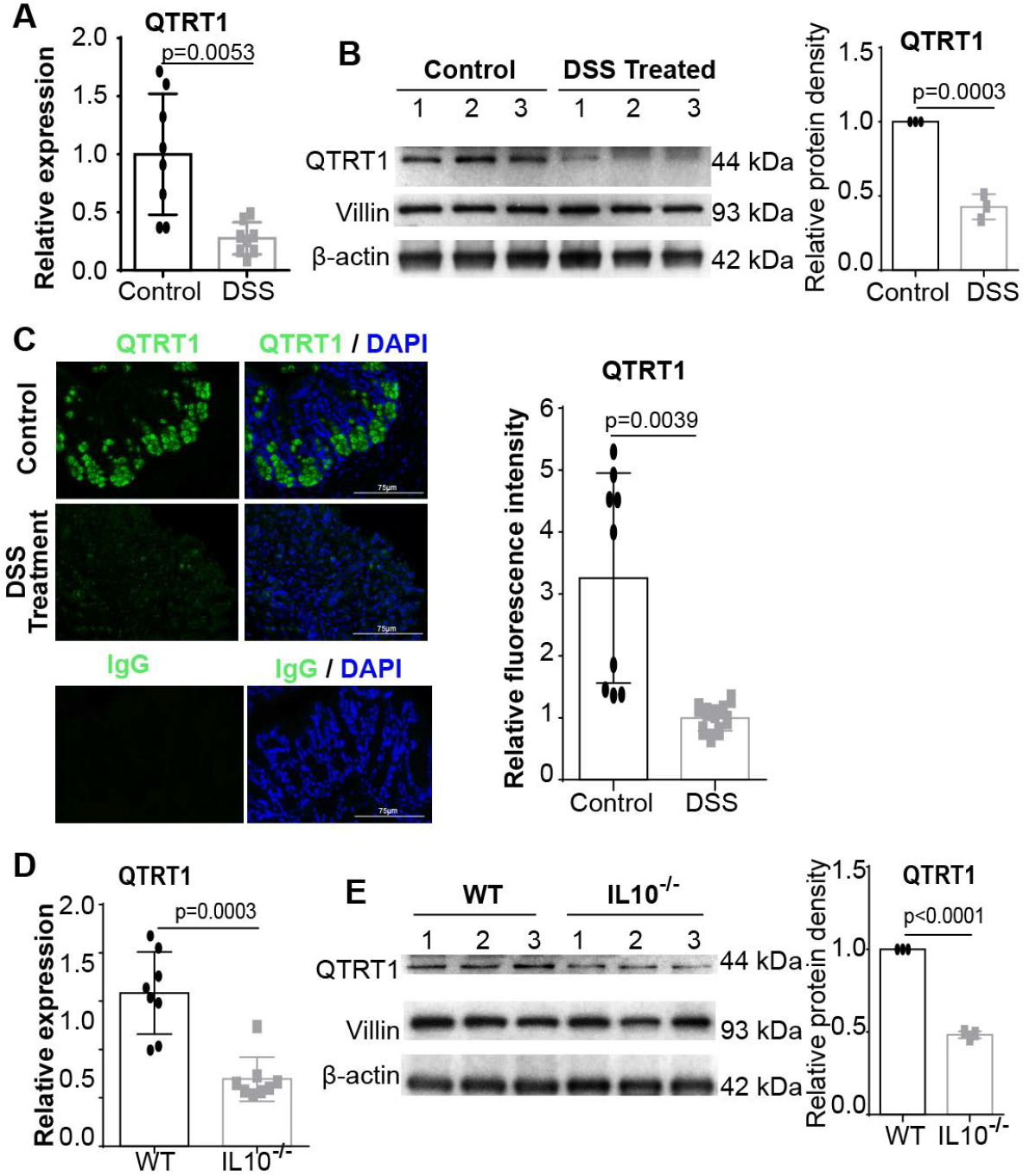
Expression of QTRT1 and junction proteins in the IBD mouse model. (**A**) mRNA expression of QTRT1 in DSS (dextran sulfate sodium)-reduced colitis mice. Data are expressed as the mean ± SD, Welch’s t test. n=4 (one-repeat well) mice per group. (**B**) QTRT1 protein expression levels were evaluated in the intestinal epithelial cells of mice treated with dextran sulfate sodium (DSS) and controls using western blotting. Data are expressed as the mean ± SD, Welch’s t test. n=3 mice per group. (**C**) QTRT1 expression in colons from DSS-treated mice was visualized by immunofluorescence staining. Mouse IgG was used as the negative experimental control in all the staining. Scale bar is 75 μm. The relative fluorescence intensity was quantified with ImageJ by counting 3-image each sample. Data are shown as the mean ± SD, Welch’s t test. n=3 per group. (**D**) mRNA expression of QTRT1 in IL-10 knockout mice. Data are expressed as the mean ± SD, Welch’s t test. n=4 (one-repeat well) mice per group. (**E**) QTRT1 protein levels were evaluated in the intestinal epithelial cells of IL10^−/-^mice and wild-type mice (WT) using western blotting. Data are expressed as the mean ± SD, Welch’s t test. n=3 mice per group.

As one of the most commonly used IBD models, IL-10-deficient mice (IL-10^−/-^) spontaneously develop chronic enterocolitis ^21, 22^. This spontaneous onset of gut inflammation (colitis) in IL-10-deficient mice is characterized by histological findings that are similar to those of human IBD ^23^. Here, we further investigated the Q-tRNA modification, mainly QTRT1 expression, in the colon of IL-10-deficient mice with inflammation. After colonic epithelial cell collection and total RNA extraction, we detected the mRNA level of QTRT1 using real-time PCR. Similar to the results we found in DSS-induced colitis, QTRT1 expression in IL-10^−/-^ mice was significantly suppressed compared with that in wild-type (WT) mice (p<0.001) (Figure 2D). Then, we further performed western blot analysis to evaluate the protein expression level. We detected significantly suppressed expression of QTRT1 in IL-10^−/-^ mice compared with WT mice (p<0.001) (Figure 2E). Taken together, we found significantly reduced QTRT1 expression in DSS-induced colitis and IL10^−/-^ mouse models, suggesting dysfunction of Q-modification in the inflamed intestine.

### Reduced QTRT1 was associated with alterations in β-catenin and Claudins in colitis mouse models

To investigate the regulatory genes correlated with the downregulation of QTRT1 in intestinal inflammation, we evaluated the expression of adhesion regulators and tight junction regulators in colitis mouse models. As a multitasking and evolutionarily conserved molecule, β-catenin has a crucial role in a multitude of developmental and homeostatic processes, such as cell proliferation and adhesion ^24^. We found significantly downregulated expression of β-catenin at the mRNA level in colon tissue from DSS-treated mice compared with nontreated mice (p<0.01) (Figure 3A). Meanwhile, the protein expression of β-catenin was also significantly downregulated in the colon of DSS-treated mice (p<0.05) (Figure 3B). Importantly, we found a significantly positive correlation of the QTRT1 alteration (Figure 2) and β-catenin alteration at both mRNA (correlation=0.93, p=1.1×10^−7^) and protein level (correlation=0.91, p=0.011) using the Spearman Correlation test (Figure 3D). Furthermore, immunofluorescence staining of β-catenin was applied to colon tissue to verify the changes in β-catenin, which was significantly downregulated (fluorescence intensity) in mice with colitis (p<0.01) (Figure 3C). All these findings highlighted the critical role of Q-tRNA modification in β-catenin-related cell proliferation and colitis.

**Figure 3.**
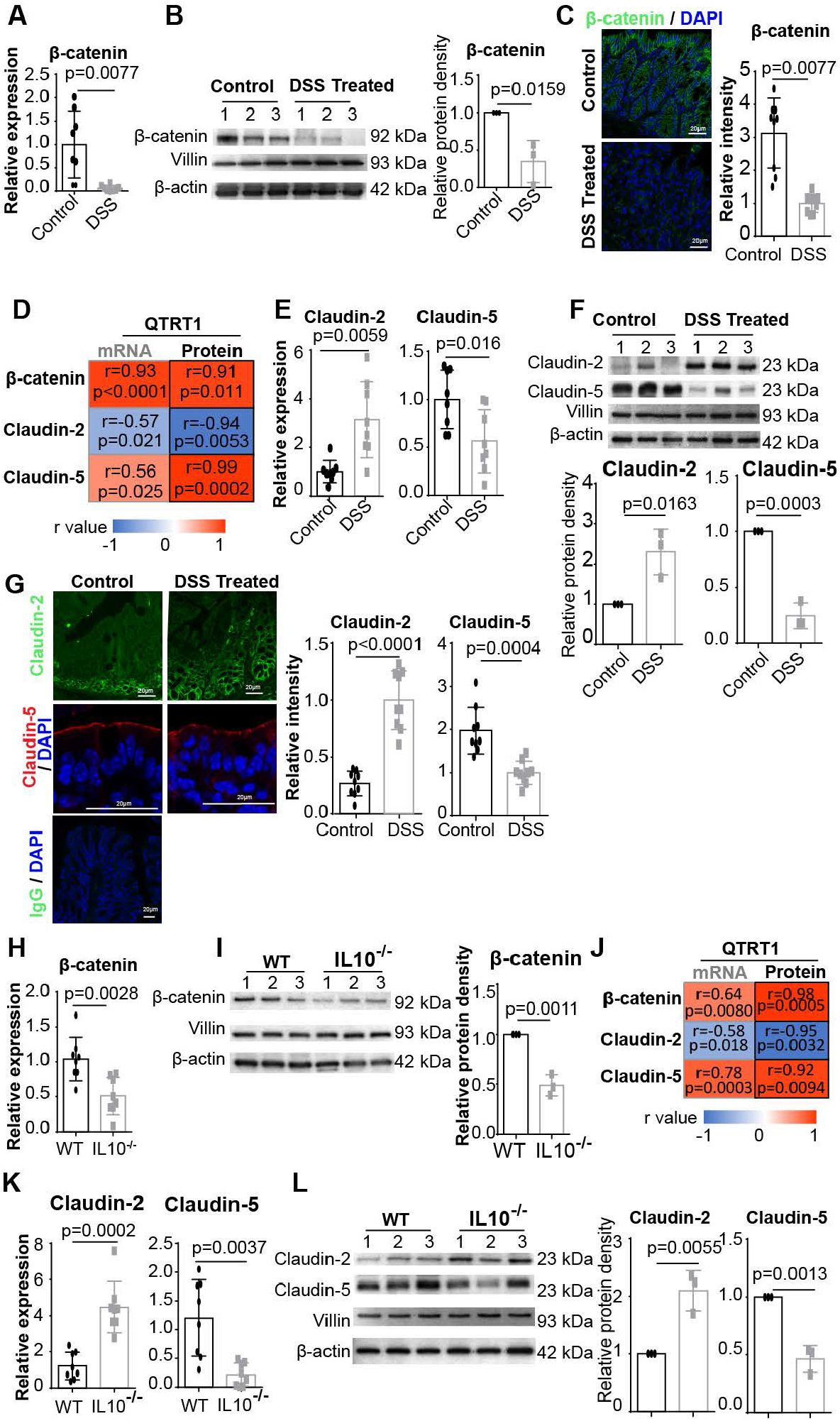
Expression of β-catenin and Claudin proteins in colitis mice and controls. (**A**) mRNA expression of β-catenin in DSS-reduced colitis mice. Data are expressed as the mean ± SD, Welch’s t test. n=4 (one-repeat well) mice per group. (**B**) β-Catenin protein expression levels were evaluated in the intestinal epithelial cells of mice treated with DSS and controls using western blotting. Data are expressed as the mean ± SD, Welch’s t test. n=3 mice per group. (**C**) β-catenin expression in colons from DSS-treated mice was visualized by immunofluorescence staining. The scale bar is 20 μm. The relative fluorescence intensity was quantified with ImageJ by counting 3-image each sample. Data are shown as the mean ± SD, Welch’s t test. n=3 per group. (**D**) The correlation of QTRT1 alteration in Fig 2. and alterations of β-catenin and Claudins was analyzed in mRNA (Real-time PCR) and protein (western blot) level using Spearman’s rank correlation coefficient. The correlation coefficient value r and p-value were indicated in the frames and color bar. (**E**) mRNA expression of Claudin-2 and Claudin-5 in DSS-induced colitis and control mice. Data are expressed as the mean ± SD, Welch’s t test. n=4 (one-repeat well) mice per group. (**F**) Claudin protein levels were evaluated in the intestinal epithelial cells of DSS-treated mice and control mice using western blotting. Data are expressed as the mean ± SD, Welch’s t test. n=3 mice per group. (**G**) Claudins in the colon from DSS-treated mice were visualized by immunofluorescence staining. The scale bar is 20 μm. The relative fluorescence intensity was quantified with ImageJ by counting 3-image each sample. Data are shown as the mean ± SD, Welch’s t test. n=3 per group. IgG was used as the negative experimental control in all IF staining. (**H**) mRNA expression of β-catenin in IL-10 knockout and wild-type mice. Data are expressed as the mean ± SD, Welch’s t test. n=4 (one-repeat well) mice per group. (**I**) β-Catenin protein expression levels were evaluated in the intestinal epithelial cells of IL-10 knockout and wild-type mice using western blotting. Data are expressed as the mean ± SD, Welch’s t test. n=3 mice per group. (**J**) The correlation of QTRT1 alteration in Fig 2. and alterations of β-catenin and Claudins was analyzed in mRNA (Real-time PCR) and protein (western blot) level using Spearman’s rank correlation coefficient. The correlation coefficient value r and p-value were indicated in the frames and color bar. (**K**) mRNA expression of Claudin-2 and Claudin-5 in IL-10 knockout and wild-type mice. Data are expressed as the mean ± SD, Welch’s t test. n=4 (one-repeat well) mice per group. (**L**) Claudin protein levels were evaluated in the intestinal epithelial cells of IL-10 knockout and wild-type mice using western blotting. Data are expressed as the mean ± SD, Welch’s t test. n=3 mice per group.

Tight junctions (TJs) are intercellular adhesion complexes in intestinal epithelia and endothelia that control paracellular permeability. Claudins are a family of transmembrane proteins that form gated ion-selective paracellular pores through the paracellular diffusion barrier, in which Claudin-5 is responsible for the watertight stability of the cell, whereas Claudin-2 is permeable to both ions and water ^25^, thus ensuring proper availability of water and ions for effective cellular functions. Here, in our DSS-induced colitis mouse model, significantly upregulated mRNA expression of Claudin-2 (p<0.01) and significantly downregulated Claudin-5 (p<0.05) were detected in the colon of DSS-treated mice compared with nontreated mice (Figure 3E). Similarly, the protein expression alteration of Claudin-2 and Claudin-5 was analyzed by western blot, in which we found alterations consistent with the mRNA alterations, of which Claudin-2 was significantly upregulated (p<0.05) while Claudin-5 was significantly downregulated (p<0.001) (Figure 3F). Using the Spearman correlation test, we found that these protein alterations in Caludin-2 (correlation= -0.94, p=0.0053) and Claudin-5 (correlation= 0.99, p=0.00017) were significantly correlated with the protein alteration of QTRT1 (Figure 2) in DSS-treated mice and control mice (Figure 3D). Moreover, we further verified the alterations of Claudin-2 and Claudin-5 using immunofluorescence staining and found that the fluorescence was significantly upregulated for Claudin-2 (p<0.001) and downregulated for Claudin-5 (p<0.001) (Figure 3G). However, there was no fluorescence staining in the IgG negative control used in the whole process of all IF staining described in this section. Furthermore, we found that these alterations were correlated with QTRT1 downregulation.

We then investigated the correlation of QTRT1 regulation and cell proliferation and intestinal junction functions in IL-10-deficient mice. First, we evaluated the mRNA expression of β-catenin using real-time PCR, in which we found significant downregulation of β-catenin compared with that in wild-type mice (p<0.05) (Figure 3H). These changes in β-catenin were also verified at the protein level by western blot analysis using IL-10-deficient mice and wild-type mice (p<0.001) (Figure 3I). Moreover, these protein alterations of β-catenin in IL-10 deficient mice significantly correlated with the downregulation of QTRT1 expression (Figure 2) in both mRNA (correlation= 0.64, p=0.0080) and protein level (correlation= 0.98, p=0.00050) using Spearman Correlation test (Figure 3J). Then, we evaluated the mRNA expression of Claudin-2 and Claudin-5 in these IL-10-deficient mice using real-time PCR as described above. Similar to the results we found in DSS-induced colitis, significantly upregulated mRNA expression of Claudin-2 (p<0.01) and downregulated Claudin-5 (p<0.01) were found in IL-10-deficient mice (Figure 3K). Using western blotting, we then investigated the protein expression levels of the target genes and found significantly upregulated Claudin-2 (p<0.01) and downregulated Claudin-5 (p<0.01) in colon tissue from IL-10-deficient mice compared with wild-type mice (Figure 3L). Moreover, we found these alterations of Claudin-2/Claudin-5 were significantly correlated to the changed QTRT1 in both mRNA (Claudin-2: correlation=-0.58, p=0.018; Claudin-5: correlation=0.78, p=0.00034) and protein (Claudin-2: correlation=-0.95, p=0.0032; Claudin-5: correlation=0.92, p=0.0094) level (Figure 3J). Taken together, we found the downregulation of β-catenin and Claudin-5 and the upregulation of Claudin-2 in the colon from intestinal inflamed mice (DSS treatment and IL-10-deficient), and these alterations were correlated with QTRT1 downregulation (Figure 3). These findings highlighted the importance of QTRT1 in cell proliferation and intestinal barrier function *in vivo*.

### Reduced QTRT1 was associated with alteration of adhesion and junctions in vitro

Because of the critical role of QTRT1 in the inflamed intestine in our in vivo mouse model, we then investigated the impact of QTRT1 suppression on cell function in vitro. Using a loss-of-function study, we suppressed the expression of QTRT1 to investigate its regulatory role in cell proliferation and junction proteins. To this end, we suppressed the expression of QTRT1 in Caco2 BBE cells using the double nickase plasmid, which was indicated by the significantly downregulated QTRT1 expression at both the mRNA and protein levels (Figure 4). The cell proliferation ability was significantly suppressed by the knockdown (KD) of QTRT1 in Caco2 BBE cells, which was indicated by the downregulated expression of proliferating cell nuclear antigen (PCNA) in the knockdown cells, compared with the controls (Figure 4A and B). Meanwhile, the expression of adhesion and tight junction proteins, including β-catenin, Claudin-2 and Claudin-5, was also changed in QTRT1-KD cells, which is consistent with the findings in vivo. Using real-time PCR, western blotting and immunofluorescence staining, we found that the expression of β-catenin and Claudin-5 was significantly downregulated in QTRT1-KD cells compared with control Caco2 BBE cells at both the mRNA (p<0.05) (Figure 4A) and protein levels (p<0.001) (Figure 4B and C). However, the knockdown of QTRT1 significantly upregulated the expression of Claudin-2 in QTRT1-KD Caco2 BBE cells at both the mRNA (p<0.01) (Figure 4A) and protein levels (p<0.01) (Figure 4B). There is no fluorescence staining in the IgG negative control used in the whole process of the IF staining described in this section.

**Figure 4.**
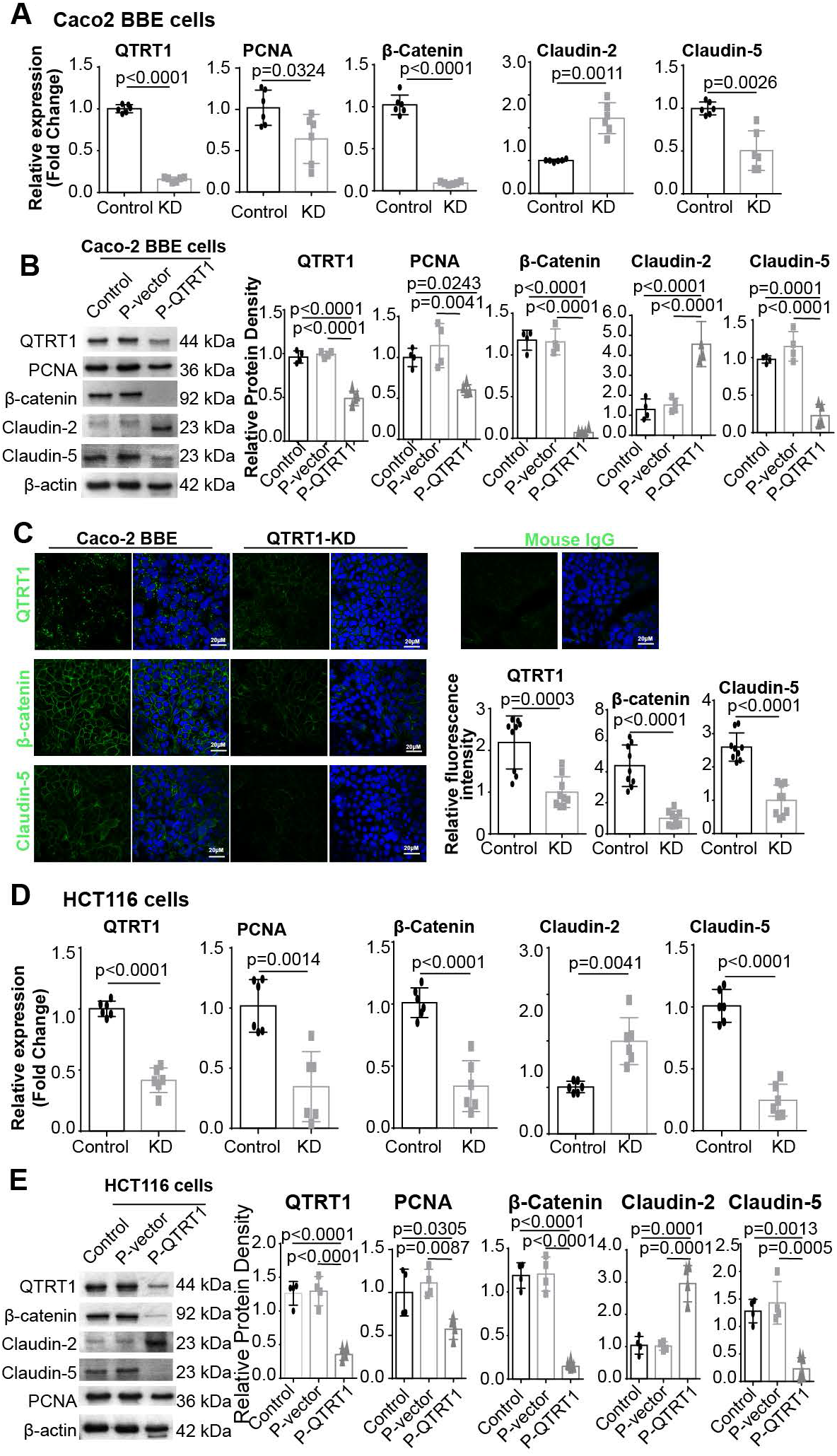
Expression of β-catenin and Claudins in QTRT1-knockdown cells. (**A**) The mRNA expression levels of targeted genes in Caco-2 BBE cells, including QTRT1 knockdown (KD) by transfection with QTRT1 Double Nickase Plasmid (h) (Santa Cruz, sc-413456-NIC) and controls (Control), were measured by real-time PCR. Data are expressed as the mean ± SD, Welch’s t test. n=3 (one-repeat well) per group. (**B**) Protein expression of targeted genes in Caco2 cells transfected with QTRT1 double nickase plasmid (h) or control double nickase plasmids (P-vector) and negative control (Control) was evaluated by western blotting. Data are expressed as the mean ± SD, one-way ANOVA. n=4 per group. (**C**) Targeted protein expression in QTRT1 knockdown Caco-2 BBE cells was visualized by immunofluorescence staining. The scale bar is 20 μm. The relative fluorescence intensity was quantified with ImageJ by counting 3-image each sample. Data are shown as the mean ± SD, Welch’s t test. n=3 per group. (**D**) The mRNA expression levels of targeted genes in HCT116 cells, including QTRT1 knockdown (KD) by transfection with QTRT1 double nickase plasmid (h) and controls (control), were measured by real-time PCR. Data are expressed as the mean ± SD, Welch’s t test. n=3 (one-repeat well) per group. (**E**) Protein expression of targeted genes in CHT116 cells transfected with QTRT1 double nickase plasmid (h) or P-vector and negative control (control) was evaluated by western blotting. Data are expressed as the mean ± SD, one-way ANOVA. n=4 per group.

To further verify our findings of the influences of QTRT1 knockdown on cell proliferation and junction functions, we evaluated the mRNA and protein expression of β-catenin and Claudins in human HCT116 colon epithelial cells. The significantly downregulated mRNA expression (Figure 4D) and protein expression (Figure 4E) of QTRT1 in the HCT116-KD cells clearly showed the knockdown of QTRT1 in these cells. Cell proliferation was suppressed after knockdown of QTRT1, as indicated by the downregulated expression of PCNA (Figure 4D and E). Similarly, we found the downregulation of β-catenin and Claudin-5 and the upregulation of Claudin-2 in QTRT1-KD HCT116 cells, compared with control cells (Figure 4D and E). Taken together, these findings clearly showed the impact of QTRT1 knockdown on cell proliferation and junctions by altering the expression of PCNA, β-catenin and claudins in two human intestinal epithelial cell lines.

### Alteration of QTRT1-related metabolites

The guanine is exchanged from the tRNA-containing guanosine with queuine to obtain queuosine-containing tRNA (Q-tRNA), whose reaction is controlled by the eukaryotic tRNA-guanine transglycosylase (eTGT)-composed subunits of QTRT1 and QTRT2. Therefore, the dysfunction of QTRT1, the catalytic subunit of eTGT, in IBD patients may further influence the abundance of metabolites related to QTRT1-included pathways. Thereafter, the abundance of guanine in the feces was compared between IBD patients and controls by revisiting metabolite data from the Metabolomics Workbench (https://www.metabolomicsworkbench.org/).

There are two datasets of IBD research: the project PR000639 with 145 UC patients, 266 CD patients and 134 controls without any IBD symptoms ^26^, and the project PR000677 with 76 UC patients, 88 CD patients, and 56 controls without any IBD symptoms ^27^. For the UC patients, the abundance of guanine was significantly downregulated compared with controls in both projects (p<0.001 and p<0.05, respectively) (Figure 5A). Similarly, guanine abundance in the feces of patients with CD symptoms was significantly downregulated compared with controls without CD symptoms in both projects used in this study (p<0.001 and p<0.05, respectively) (Figure 5B).

**Figure 5.**
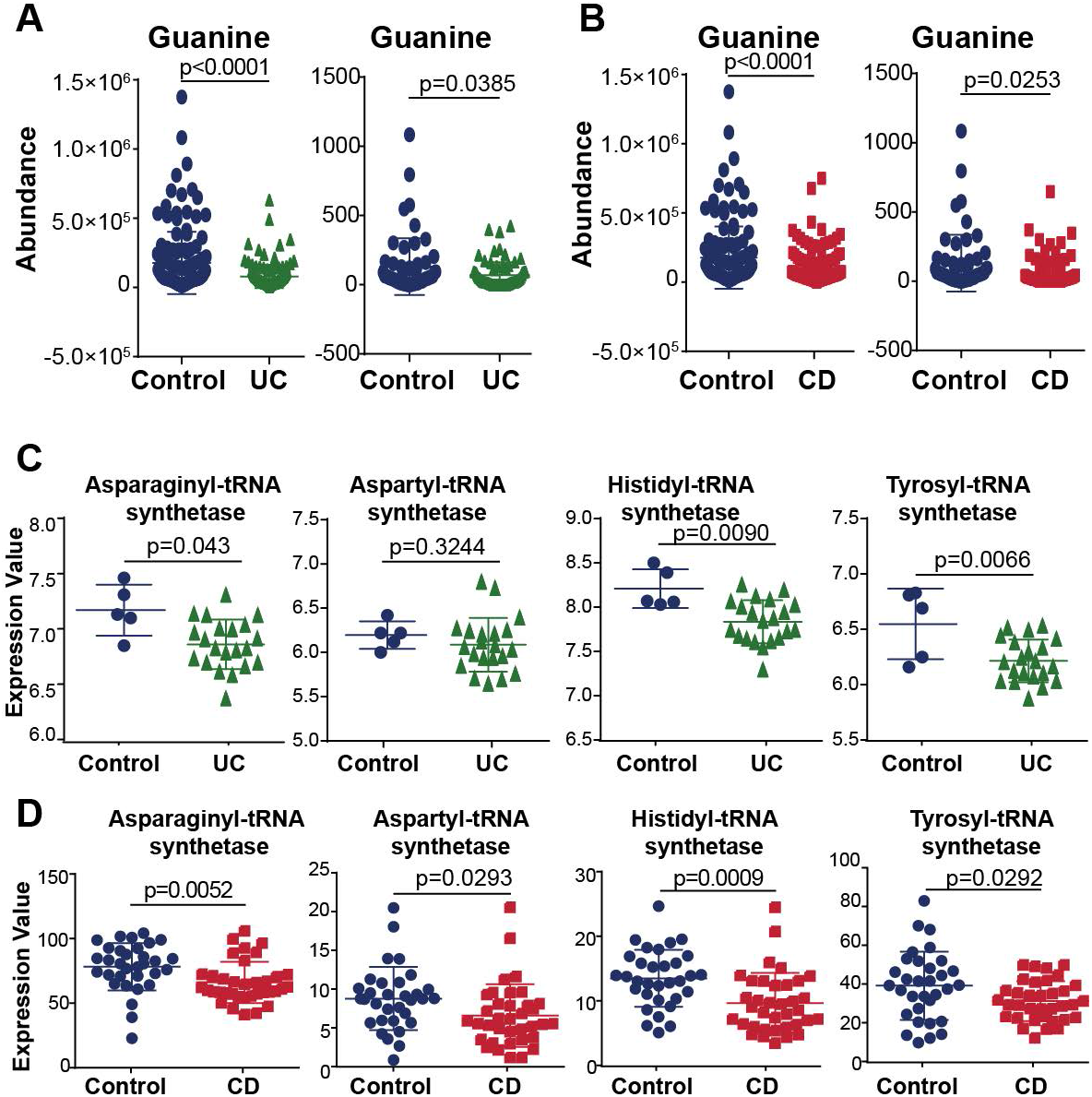
Alterations of metabolites related to QTRT1 in human IBD patients. The guanine is exchanged from the tRNA-containing guanosine with queuine to obtain queuosine-containing tRNA, whose reaction is controlled by the eukaryotic tRNA-guanine transglycosylase (eTGT)-composed subunits of QTRT1 and QTRTD1. The abundance of guanine was compared between IBD patients and controls by revisiting metabolite data from Metabolomics Workbench (https://www.metabolomicsworkbench.org/). Alteration of guanine in UC (**A**) and CD (**B**) patients. Two datasets were included in the analysis with project numbers PR000639 (UC: n=145; CD: n=266; control: n=134) and PR000677 (UC: n=76; CD: n=88; control: n=56). The data are shown as the mean ± SD, Welch’s t test. (**C**) Four synthetases of tRNAs (asparaginyl, aspartyl, histidyl, and tyrosyl) associated with queuine metabolism in eukaryotic cells were compared between UC patients (n=14) and controls (n=5) sourced from GEO datasets with accession number GSE9452. (**D**) Four synthetases of tRNAs (asparaginyl, aspartyl, histidyl, tyrosyl) were compared between CD patients (n=36) and controls (n=32) sourced from ArrayExpress (https://www.ebi.ac.uk/arrayexpress) with accession number E-MTAB-5783. The data are shown as the mean ± SD, Welch’s t test.

In eubacteria and eukaryotes, queuine is found as the sugar nucleotide queuosine within the anticodon loop of transfer RNA isoacceptors for the amino acids tyrosine, asparagine, aspartic acid and histidine ^1^. Because of the dysfunction of QTRT1 and alteration of guanine in IBD patients, it is reasonable to hypothesize that the changed Q-tRNA in IBD patients further alters the related tRNA synthetases. Therefore, we investigated the synthetase expression level of these four tRNAs, which are related to Q-tRNA modification, in both UC and CD patients and normal controls from the available datasets. We found that three tRNA synthetases, asparaginyl-tRNA synthetase (p<0.05), histidyl-tRNA (p<0.01) and tyrosyl-tRNA synthetase (p<0.01), were significantly downregulated in the intestinal mucosa of UC patients (n=14) compared with controls (n=5) from GSE9442 (Figure 5C). Although there were no significant differences in aspartyl-tRNA synthetase between UC patients and controls (p=0.32), a trend of downregulation was observed (Figure 5C). Meanwhile, all four rRNA synthetases, including asparaginyl tRNA synthetase (p<0.01), aspartyl-tRNA synthetase (p<0.05), histidyl-tRNA (p<0.001) and tyrosyl-tRNA synthetase (p<0.05), were significantly downregulated in the intestinal mucosa of CD patients (n=36) compared with controls (n=32) (Figure 5D).

### Queuine treatment enhances the proliferation and tight junctions of epithelial cells in vitro

To further evaluate the critical role of QTRT1 and related molecules in cell biosynthesis, we treated Caco-2 BBE cells with q, which is the substrate of the biosynthesis of Q-tRNA. As we found above that QTRT1 deletion markedly impacted cell proliferation, we measured the cell proliferation activity of the cells using the MTT assay. The cell proliferation activity was significantly (p<0.05) upregulated after 24 hours of treatment with queuine, compared with that in untreated cells. This upregulation lasted from 48 hours to 72 hours post-treatment (Figure 6A). This upregulated cell proliferation was further verified by the increased expression of the cell proliferation marker PCNA using real-time PCR (p<0.05) (Figure 6B). We also verified this upregulation of PCNA using western blot analysis, which showed significantly higher expression compared with controls (p<0.01) (Figure 6C). Using immunofluorescence staining of Ki67, another cell proliferation marker, we found a significantly higher Ki67-positive cell ratio in queuine-treated Caco2 BBE cells than in controls (p<0.01) (Figure 6D), indicating high cell proliferation activities in queuine-treated cells.

**Figure 6.**
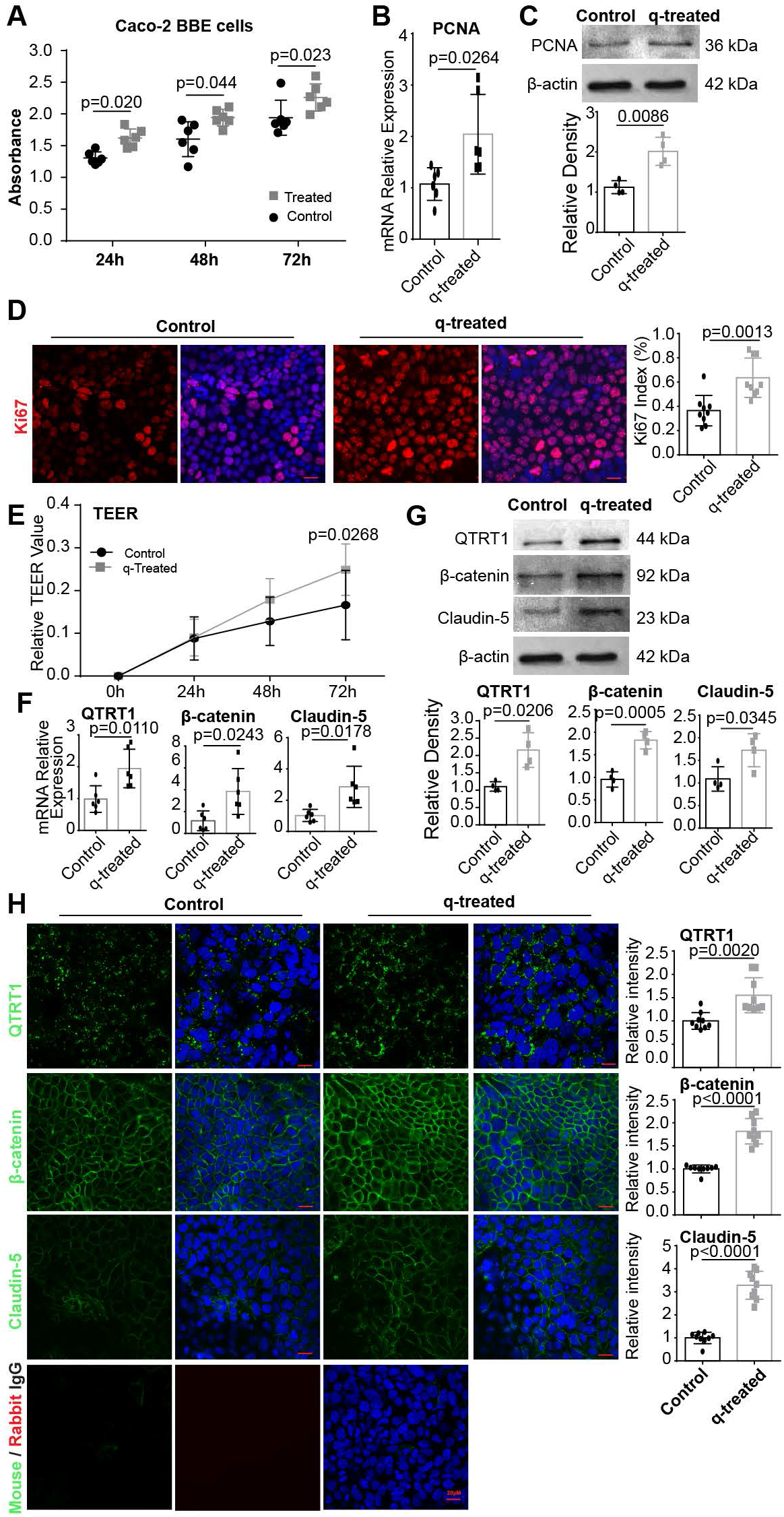
Queuine treatment regulates cell proliferation and permeability. (**A**) An MTT assay showing the proliferation of control and Caco-2 BBE cells treated with queuine hydrochloride was conducted at 24 hours, 48 hours and 72 hours after treatment. Data are expressed as the mean ± SD, generalized linear mixed models. n=6 per group. (**B**) The mRNA expression levels of PCNA in Caco-2 BBE cells treated with queuine for 72 hours were measured by real-time PCR. Data are expressed as the mean ± SD, Welch’s t test. n=3 (one-repeat well) per group. (**C**) The protein expression of PCNA in Caco-2 BBE cells treated with queuine for 72 hours was measured by western blotting. Data are expressed as the mean ± SD, Welch’s t test. n=4 per group. (**D**) Immunofluorescence staining of the cell proliferation marker Ki67 was performed in Caco-2 BBE cells after 72 hours of queuine treatment. The Ki67 index was calculated with the formula: Ki67-positive cells/total cells. The relative fluorescence intensity was quantified with ImageJ by counting 3-image each sample. Data are expressed as the mean ± SD, Welch’s t test. n=3 per group. (**E**) The TEER value was measured and monitored on HT-29 cells at 24-hour, 48-hour and 72-hour after treating with queuine. Data was expressed as mean ± SD, two-way ANOVA. n=6 per group. (**F**) The mRNA expression levels of QTRT1, β-catenin, and Claudin-5 in Caco-2 BBE cells treated with queuine for 72 hours were measured by real-time PCR. Data are expressed as the mean ± SD, Welch’s t test. n=3 (one-repeat well) per group. (**G**) The protein expression of QTRT1, β-catenin and Claudin-5 in Caco-2 BBE cells treated with queuine for 72 hours was measured by western blotting. Data are expressed as the mean ± SD, Welch’s t test. n=3 per group. (**H**) Immunofluorescence staining of the cell proliferation markers QTRT1, β-catenin and Claudin-5 was performed in Caco-2 BBE cells after 72 hours of queuine treatment. The relative fluorescence intensity was quantified with ImageJ by counting 3-image each sample. Data are expressed as the mean ± SD, Welch’s t test. n=3 per group.

Then, we investigated the impact of queuine treatment on cell permeability in vitro. Using the TEER assay, we found a significantly higher TEER value after treating the cells for 72 hours with queuine (p<0.05) (Figure 6E). These altered cell permeability and cell proliferation benefited from the upregulated expression of QTRT1 in the q-treated cells, which was indicated at both the mRNA level by real-time PCR (p<0.05) (Figure 6F) and the protein level by western blot (p<0.05) (Figure 6G), which indicated the enhanced function of QTRT1-containing eTGT in this process. This upregulation of QTRT1 in queuine-treated Caco2 BBE cells was also detected using immunofluorescence staining (Figure 6H). As an important functional protein in cell permeability, alterations in β-catenin and Claudin-5 were evaluated in queuine-treated Caco2 BBE cells. As shown in Figure 6F, the mRNA expression of β-catenin (p<0.05) and Claudin-5 (p<0.05) was significantly upregulated after treating the cells with queuine compared with untreated cells. Similarly, these alterations were further verified by western blot (Figure 6G) and immunofluorescence staining (Figure 6H). There was no fluorescence staining in the IgG negative control used in the whole process of IF staining.

We used human colonoids to further investigate the benefits of queuine treatment. After treating the colonoids for 96 hours, the relative organoid volume size was significantly increased in the treated group compared with the control group (p<0.05) (Figure 7A). These findings clearly showed the functional role of queuine in alterations of intestinal cell proliferation. However, more studies need to be performed to investigate the potential underlying mechanisms. Taken together, these results clearly showed the importance of queuine and QTRT1 in cell proliferation and cell permeability, suggesting the potential treatment strategy for IBD patients using queuine or queuine-related compounds.

**Figure 7.**
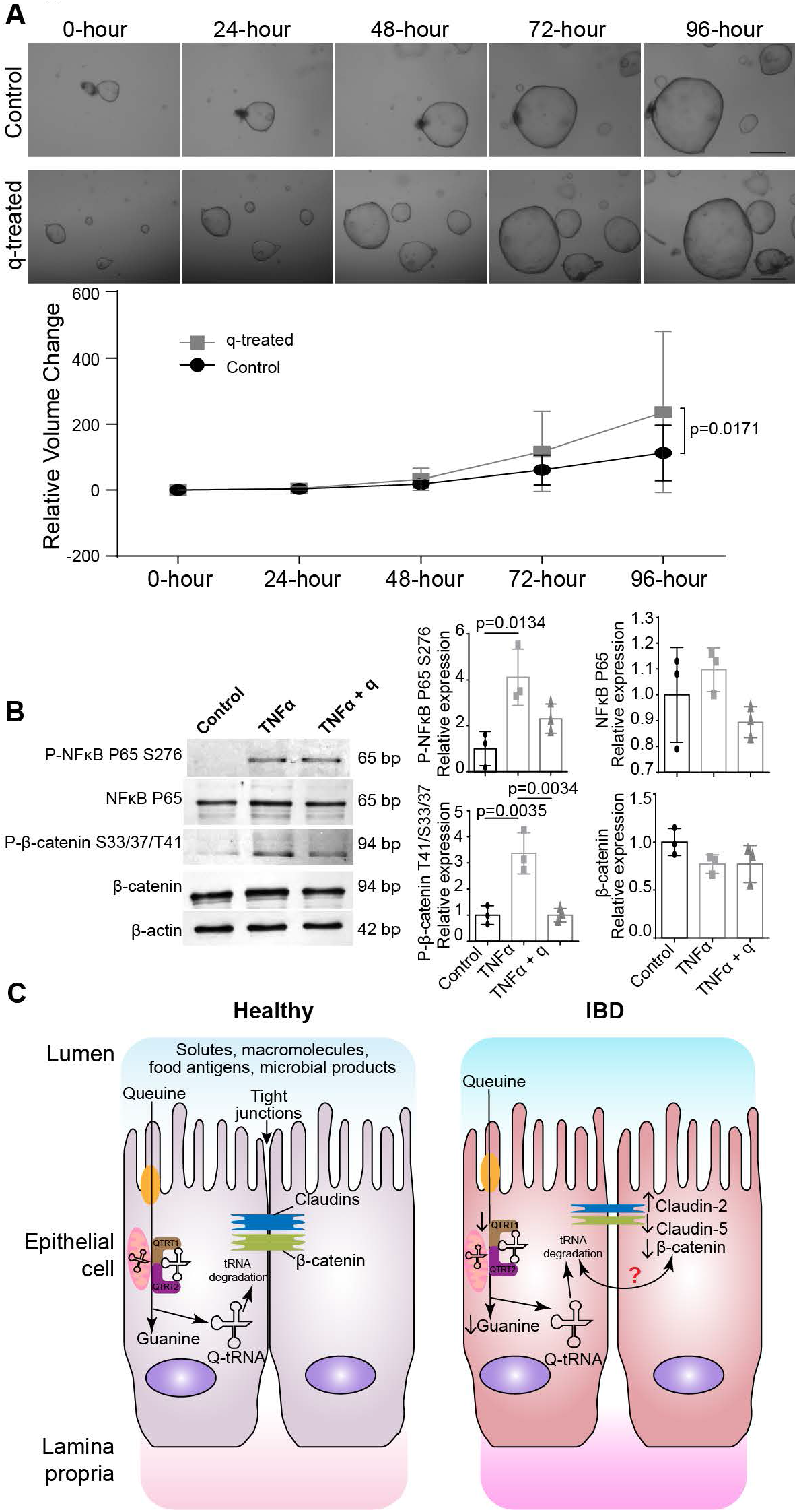
Queuine treatment in human colonoids. (**A**) Human colon organoids from healthy people were treated with queuine hydrochloride and monitored for 96 hours. The volume of the organoids was calculated with the formula: volume (μm^3^) = [(4 × π × (dimeter/2)^2^]/3. The scale bar is 750 μm. Data are expressed as the mean ± SD, two-way ANOVA. n=10 per group. (**B**) Human colonic HT29 cells were treated with TNF-α for 1 hour. Data are expressed as the mean ± SD, one-way ANOVA. n=3 per group. (**C)** The working model of this study. Queuine is produced by bacteria and salvaged by human body via intestinal epithelium. Queuine replaces guanine at the wobble position of tRNAs and promotes efficient mRNA translation. The gut microbial dysbiosis, one of them most common symptoms of the IBD patients, could critically impact on the functions of intestinal epithelium and queuine-to-guanine transportation, and mRNA translation. The protein expression of QTRT1, one of the subunits of the queuine-insertase complex, is suppressed in IBD patients. Reduced QTRT1 led to increased permeability and dysfunction of intestinal epithelial cells through Claudin-5 and β-catenin.

### Queuine treatment reduced TNF-α-induced inflammation in vitro

We further investigated the protective role of queuine in inflammation. In HT29 cells treated with TNF-α, we found that phosphorylated NF-κB P65 S276 and phosphorylated β-Catenin S33/37/T41 were increased (p<0.05). Queuine reduced the expression of p-P65 and p-β-Catenin (S33/37/T41) in TNF-α/queuine-treated cells compared to TNF-α-treated cells (p<0.05) (Figure 7B). These findings indicated the protective role of queuine in intestinal epithelial inflammation.

## Discussion

In the current study, we reported that QTRT1 is significantly downregulated in human IBD patients, including UC and CD patients. In two IBD mouse models, the reduction in QTRT1 was significantly correlated with cell proliferation and intestinal barrier functions through altered Claudin-2, Claudin-5, and β-catenin. Queuine is able to enhance cell proliferation and intestinal barriers in cell cultures and human colonoids. Moreover, the downstream metabolite guanine of QTRT1 was involved in the process, and QTRT1-related tRNA synthesis was markedly altered in IBD patients.

Q-tRNA modification is critical for fidelity and accuracy when translating RNA to protein. Dysfunction of Q-tRNA is associated with cancer proliferation and malignancy ^14, 28, 29^. However, the mechanisms by which Q and Q-tRNA modifications influence intestinal epithelial biology and chronic inflammation are unknown. For the first time, our study demonstrated the downregulation of QTRT1 and Q-tRNA dysfunction in IBD. Furthermore, we provided evidence of altered downstream molecules, including guanine and four tRNA synthetases, of QTRT1-containing enzyme catalysis in human IBD. In eukaryotes, queuine is the precursor base to the queuosine tRNA modification, which is processed based on the enzyme complex containing QTRT1 subunits ^1, 30^. In eubacteria and eukaryotes, guanine is exchanged from tRNA containing guanosine with queuine to obtain Q-tRNA, the reaction of which is controlled by eukaryotic tRNA-guanine transglycosylase (eTGT), which is composed of QTRT1 and QTRT2 ^1, 30^. Q-tRNA deficiency was found to be correlated with a variety of diseases, such as tumor growth, encephalomyelitis, and leukemia ^1, 14, 31, 32^. Dysfunction of the eukaryotic tRNA-guanine transglycosylase complex, which contains QTRT1, is reported to be involved in cancer proliferation and malignancy ^14^. Our studies on t-RNA regulation of β-catenin, a proliferation regulator, further supported the importance of Q-tRNA in health and disease.

We reported that QTRT1 downregulation was related to altered β-catenin, Claudin-2, and Claudin-5. It is well known that TJs are essential to the function of the physical intestinal barrier by regulating the paracellular movement of various substances and waste across the intestinal epithelium. TJ dysfunctions are reported in IBD and other inflammatory diseases ^33-35^. Meanwhile, being the link between amino acids and nucleic acids, tRNA functions as an ‘adaptor’ molecule that translates three-nucleotide codon sequence in the mRNA into the suitable amino acid of that codon ^36^. It is reasonable to hypothesize that the depleted expression of QTRT1 impacts Q-tRNA modification, which may further influence tRNA on amino acid transferring. Thereafter, inhibited tRNA activity impacted mRNA transcription and protein translation. The QTRT1 regulation of β-catenin and Claudin-2/5 was further confirmed in cells with QTRT1 knockdown. On the one hand, QTRT1 knockout led to decreased protein levels of β-catenin and Claudin-5 and increased Claudin-2, suggesting decreased cell adhesion and increased permeability among cells. On the other hand, when the cells were treated with queuine, the substrate of the QTRT1-containing enzyme complex, the cell proliferation activity and Claudin-5 expression were upregulated. These data indicated a novel role of QTRT1 in regulating intestinal epithelial cell junctions.

The sugar nucleotide queuosine within the anticodon loop of transfer RNA is an acceptor for the amino acids tyrosine, asparagine, aspartic acid, and histidine ^1, 30^. Our studies provided evidence that lower mRNA expression levels of tRNAs transfer these four amino acids in IBD patients than in healthy controls. We also found that the abundance of guanine was significantly lower in UC and CD patients. These findings indicated the alteration of QTRT1-related metabolites and tRNA synthetases in IBD patients, which highlighted the critical role of QTRT1 in the biosynthesis of Q-tRNA and metabolites in the related pathways.

Queuine is an elusive and less-recognized micronutrient acquired from the diet or microbiome ^1^. Micronutrients are essential elements needed by life in small quantities. They include microminerals and vitamins. Micronutrients from the diet and microbiota are essential to human health. Queuine is a microbiome/diet-derived chemical incorporated into the wobble position of tRNAs to affect fidelity and efficiency of translation from RNA to proteins. It is synthesized *de novo* in bacteria; mammals acquire queuine as micronutrient from their diet or intestinal microflora. Q-tRNA modification levels are highly dynamic and reflect the interplay between the host and microbiome. Understanding the cellular and organismal mechanisms of this microbiome-dependent micronutrient will advance the prevention of IBD and improve the quality of life of patients with IBD.

In summary, our data demonstrate that QTRT1 plays a novel role in human IBD by altering intestinal cell proliferation and junctions. Studies are needed to better understand human diseases caused by aberrations in tRNA modifications, also called tRNA modopathies ^37^. The microbiome-dependent Q supply can be altered during disease development or when a limited variety of food types are ingested. Investigations on gut tRNA modification in human IBD will uncover novel molecular mechanisms for potential prevention and therapy.

