## Supplementary material for "Dysfunction of transfer RNA modifications in inflammatory bowel disease": Methods

#### Methods and Materials

##### Human IBD Datasets reanalysis

For RNA expression analyses, we took advantage of microarray data reported in the GEO (Gene Expression Omnibus) repository (https://www.ncbi.nlm.nih.gov/geo/). In particularly, gene expression data of biopsy from human intestinal mucosa were obtained from the dataset reported on Ulcerative Colitis (UC) (GEO accession number: GSE9452 [1]) and Crohn's Disease (CD) (GEO accession number: GSE83448 [2]), and normal ileum from control individuals from control individuals [3]. Four synthetases of tRNAs (asparaginyl, aspartyl, histidyl, tyrosyl) were compared between UC patients (n=14) and controls (n=5) sourced from GEO datasets (accession number GSE9452). Similarly, the four synthetases of tRNAs were compared between CD patients (n=36) and controls (n=32) sourced from ArrayExpress (https://www.ebi.ac.uk/arrayexpress) with accession number E-MTAB-5783 [4]. For metabolic compounds analyses, we revisited the datasets available in Metabolomics Workbench (https://www.metabolomicsworkbench.org/). In particularly, metabolites analysis assessed on human stool samples were obtained from the dataset reported on IBD patients and control individuals with project number PR000639 [5] and PR000677 [6]. These two datasets were performed on both UC and CD patients, project PR000639 with UC: n=145, CD: n=266, and control: n=134, while project PR000677 with UC: n=76, CD: n=88, and control: n=56.

##### Animals and animal models

Wildtype C57BL/6 mice and IL10 knockout (IL $10^{-/-}$) mice were purchase from Jackson Laboratory (Bar Harbor, Maine, USA) previously described [7]. Experiments were performed on 8-12 weeks old mice. Animals were provided the same and consistent conditions, including diet and water ad libitum and 12 h dark/light cycle, throughout the experiments. Animals were housed in the Biologic

Resources Laboratory (BRL) at the University of Illinois at Chicago and utilized in accordance with the UIC Animal Care Committee, the Office of Animal Care and Institutional Biosafety (OACIB) guidelines, and the animal protocol (number ACC 18-179). Mouse colitis model was induced by administering the wildtype C57BL/6 mice (Jackson Laboratory, Bar Harbor, Maine, USA) with 5% DSS (dextran sulfate sodium) (USB Corp. Cleveland, OH) dissolved in filter-purified and sterilized water ad libitum for 7 days, as previously described [7].

IL-10 deficiency mouse colonic samples were harvest by scraping the tissue (intestinal epithelial cells) from the colon. The whole colon tissue from the DSS treated mice was used without scraping due to the severe damage of the intestine tissue. The collected cells or tissues were kept and sonicated in lysis buffer with 1% Triton X-100, 150 mM NaCl, 10 mM Tris, pH 7.4, 1 mM EDTA, 1 mM EGTA, pH 8.0, 0.2 mM sodium ortho-vanadate and protease inhibitor cocktail, as previously described [7]. The protein concentration was measured using the BioRad Reagent (BioRad, Hercules, CA, USA) according to the products' instruction. Finally, the protein was mixed with loading buffer (50 mM Tris, pH 6.8, 100 mM dithiothreitol, 2% SDS, 0.1% bromophenol blue, and 10% glycerol) and kept in freezer until using for western blot.

**QTRT1 knockdown in the human cells**

The human cells HCT116 and Caco-2 BBE were culture in the 6-well tissue culture plate with the DMEM growth medium to 70%-80% confluence [8]. Then, the cells were transfected with 2 µg of QTRT1 Double Nickase Plasmid (sc-413456-NIC) or Control Double Nickase Plasmid (sc-437281) (Santa Cruz, Dallas, TX, USA), 10 µL LTX Lipofectamine (Thermo Scientific, Rockford, IL, USA), and 2.5 µL PLUS Reagent (Invitrogen) per well (manufacturer's protocol). QTRT1 Double Nickase Plasmid-derived GFP marker was used for positive selection of transfected cells through flow cytometry with MoFlo Astrios cell sorter (Beckman Coulter, Indianapolis, IN, USA). After selection, cells were collected and divided into 24-well plate for culturing.

**Western blot analysis**

Animal intestinal epithelial tissue was lysed in lysis buffer as described above. Cultured cells were rinsed twice in ice-cold HBSS (Hanks' balanced salt solution) (Sigma-Aldrich, Saint Louis, MO, USA) and lysed in protein loading buffer then followed by sonication (Branson Sonifier, Danbury, CT, USA) and centrifugation [8, 9]. The target proteins were detected by special primary antibody (1:1000) followed by secondary antibody conjugated to horseradish peroxidase at 1:5000 dilution. The blots were visualized by ECL chemiluminescence (Thermo Scientific, Rockford, IL, USA). All experiments were performed 3-5 times. Western blot bands were quantified using image analyzer (ImageJ, NIH, Bethesda, MD, USA). The QTRT1 and Villin monoclonal antibody was purchased from Santa Cruz Biotechnology (Dallas, TX, USA). Monoclonal antibodies of β-catenin from BD Transduction (San Jose, CA, USA) and β-actin from Sigma-Aldrich (St. Louis, MO, USA) were used in this study. Claudin-5, Claudin-2 and Claudin-7 monoclonal antibodies were purchased from Thermo Fisher Scientific (Rockford, IL, USA). All chemicals were purchased from Sigma-Aldrich unless otherwise stated.

**Immunohistochemistry staining**

Human colon tissues from CD and UC patients and controls from non-IBD patients on paraffin-embedded sections (4 μm) from our previous studies were used in this study [10, 11]. Immunohistochemistry was performed on these paraffin-embedded sections. Briefly, the paraffin sections were baked in an oven at 56 °C for 30 min. The sides were deparaffinized and rehydrated in xylene, followed by graded ethanol washes at room temperature. Antigen retrieval was achieved by boiling the slides with sodium citrate buffer (0.01 M, pH 6.0). Sides were then incubated in hydrogen peroxide (3% $H_2O_2$ in PBS) for 10 min at room temperature, followed by incubation in 5% fetal bovine serum/PBS for 1 h. After that, the slides were incubated at 4 ℃ with primary antibody at 1:100 dilution overnight. The sections were then incubated with secondary antibodies (Jackson ImmunoResearch, West Grove, PA, USA) for 1 hour at room temperature, and VECTASTAIN ABC Kit (Vector Laboratories, Burlingame, CA, USA) for 1 hour at room temperature. After that, the tissue sections were stained with DAB Substrate Kit (Vector Laboratories, Burlingame, CA, USA) and hematoxylin (Leica Biosystems, Buffalo Grove, IL, USA), and examined with confocal microscope or EVOS M5000 (Thermo Fisher Scientific, Rockford, IL, USA). The target cells were stained with QTRT1 monoclonal antibody (Santa Cruz, Dallas, TX, USA). The IgG antibody protein (Jackson ImmunoResearch, West Grove, PA, USA) was used as negative control for all the IHC staining experiments. The Semi-quantitative analysis of immunohistochemistry staining was performed using software ImageJ Fiji as described before [12].

**Immunofluorescence staining**

Fresh intestine tissue was fixed in 10% neutral buffered formalin followed by paraffin embedding. The paraffin embedded slides (4μm) were prepared with microtome as described before [13]. Before blocking, the slides were deparaffinized by baking at 56 °C for 30 min and rehydrated by washing with xylene and graded ethanol. Cells were plated on fibronectin-coated glass coverslips and cultured to monolayer in humidified chambers with 5% $CO_2$ at 37℃. Then, the slides were blocked in 5% bovine serum albumin (BSA) with 0.1% goat serum in PBS for 1 hour at room temperature to reduce nonspecific background. After that, the slides were incubated at 4 ℃ with primary antibody at 1:100 dilution overnight. The sections were then incubated with secondary antibodies and DAPI for 1 hour at room temperature, and examined with Zeiss LSM 710 confocal microscope (ZEISS, Maple Grove, MN, USA) or EVOS M5000 (Thermo Fisher Scientific,

Rockford, IL, USA). The QTRT1 monoclonal antibody was purchased from Santa Cruz Biotechnology (Dallas, TX, USA), Claudin- 2, 5, and 7 from Thermo Fisher (Thermo Fisher Scientific, Rockford, IL, USA), and β-catenin from BD Transduction (San Jose, CA, USA). The IgG antibody protein (Jackson ImmunoResearch, West Grove, PA, USA) was used as negative control for all the IF staining experiments. The fluorescence intensity was evaluated with ImageJ [14].

**Real-Time Quantitative Polymerase Chain Reaction**

Total RNA was extracted from cultured cell monolayers or tissues from the mice using TRIzol reagent (Thermo Fisher Scientific, Rockford, IL, USA) by following the products' instruction. The total RNA from DSS-treated murine tissue was further purified via lithium chloride precipitation as described before [15]. RNA reverse-transcription was performed using the iScript complementary DNA synthesis kit (Bio-Rad Laboratories, Hercules, CA, USA) according to the manufacturer's directions. Then, the reverse-transcription complementary DNA reaction products were subjected to quantitative real-time polymerase chain reaction (PCR) using the iTaq Universal SYBR green supermix (Bio-Rad Laboratories, Hercules, CA, USA) and primers from Primer Bank (Cambridge, MA) (**Table 1**) with the MyiQ single-color real-time PCR detection system (Bio-Rad Laboratories, Hercules, CA, USA). All expression levels were normalized to β-actin levels of the same sample. The percentage expression was calculated as the ratio of the normalized value of each sample to that of the corresponding untreated control cells [16].

**Queuine treatment in cells and human colonoids**

The Caco-2 BBE cells were plated at a density of $1\times10^4$ cells per well with the 12-well cell culture plate. After 24-hour of culture, the Queuine Hydrochloride (Santa Cruz, Dallas, TX, USA) treatment was performed with 5μM final concentration for 24-hour, 48-hour, and 72-hour. Then, the cell proliferation was evaluated by the MTT Cell Proliferation Assay Kit (Thermo Fisher Scientific, Rockford, IL, USA) according to the product's instructions. For western blot and immunofluorescence staining, the cells were treated with 5μM queuine hydrochloride for 72-hour.

Human colonoids from healthy people were prepared and maintained with Matrigel in wells as described before [17]. At day 6 after passage, the colonoids were treated with 5μM queuine hydrochloride (q) and monitored by EVOS M5000 (Thermo Fisher Scientific, Rockford, IL, USA). The dimeter was measured by Image J, and the volume of the organoids was calculated with formula: volume ($\mu m^3$) = $[ (4 \times \pi \times (dimeter/2)^2] / 3$.

**TEER (transepithelial electrical resistance)**

The HT-29 cells in complete media were seeded onto the 24-transwell plate at $2.5\times10^5$ cells/200 μl [18]. After 10 days of seeding, the cells were treated with 5μM queuine hydrochloride (q), and the TEER (transepithelial electrical resistance) value was measured and monitored with EVOM3 (World Precision Instruments, Sarasota, FL, USA) to test the cell permeability.

**TNF-α treatment and queuine treatment in cells**

The HT29 cells were seeded at a density of $1\times10^4$ cells per well with the 12-well cell culture plate. In the queuine treated group, the cells were treated with 5μM queuine for 72-hour before TNF-α treatment. The TNF-α (Tumor necrosis factor alpha) (R&D Systems, Minneapolis, MN, USA) was added to the monolayers with final concentration of 50μM for 1-hour. Then, the inflammation related markers were evaluated by western blot analysis as described above. The phosphor-NF-κB p65 (Ser276) antibody and Phospho-β-Catenin (Ser33/37/Thr41) were purchased from Cell

Signaling Technology (Danvers, MA, USA), and NF-κB p65 antibody and β-catenin were from BD Transduction (San Jose, CA, USA).

**Statistical analysis**

Data shown in the bar figures were the average values from at least three independent experiments with the mean ± SD. All statistical tests were two-sided. It was considered statistically significant with p-value < 0.05. One, two, and three asterisks on the bars in the available figures indicate p-values < 0.05, < 0.01, and < 0.001, respectively. The Spearman correlation coefficient test was used to analyze the correlation between two variables (protein expression) using the Hmisc package [19]. In the MTT assay, which was tested with generalized linear mixed models, the Welch's t-test was performed for the 24-hour and 48-hour, while the Wilcoxon rank sum test was performed for 72-hour analysis. The statistical analyses were conducted by GraphPad Prism 5 (GraphPad Software, Inc., La Jolla, CA, USA).

**Table S1 Real-time PCR primers used in this study.**

| Primer | Nucleotides (5' – 3') |
|---|---|
| Human QTRT1 Forward | 5'-GAA GGG CAT CAC GAC CGA A-3' |
| Human QTRT1 Reverse | 5'-CCC GGC CTT AGA CCC AGA T-3' |
| Human β-catenin Forward | 5'-AAA ATG GCA GTG CGT TTA G-3' |
| Human β-catenin Reverse | 5'-TTT GAA GGC AGT CTG TCG TA-3' |
| Human Claudin-5 Forward | 5'-GTT TTA CGA CCC GTC TGT GC-3' |
| Human Claudin-5 Reverse | 5'-AGT GGC AGG AGA AGG TCA GC-3' |
| Human Claudin-2 Forward | 5'-ACC TGC TAC CGC CAC TCT GT-3' |
| Human Claudin-2 Reverse | 5'-CTC CCT GGC CTG CAT TAT CTC-3' |
| Human PCNA Forward | 5'-CCTGCTGGGATATTAGCTCCA-3' |
| Human PCNA Reverse | 5'-CAGCGGTAGGTGTCGAAGC-3' |
| Human β-actin Forward | 5'-CAT GTA CGT TGC TAT CCA GGC-3' |
| Human β-actin Reverse | 5'-CTC CTT AAT GTC ACG CAC GAT-3' |
| Mouse QTRT1 Forward | 5'-AAT TGG CCC CAC AAT CTG CT-3' |
| Mouse QTRT1 Reverse | 5'-CTG GGC TCA AAA GTG TCT CTT C-3' |
| Mouse β-catenin Forward | 5'-TGC TGA AGG TGC TGT CTG TC-3' |
| Mouse β-catenin Reverse | 5'-CTG CTT AGT CGC TGC ATC TG-3' |
| Mouse Claudin-5 Forward | 5'-AGG CAC GGG TAG CAC TCA CG-3' |
| Mouse Claudin-5 Reverse | 5'-CAT AGT TCT TCT TGT CGT AAT C-3' |
| Mouse Claudin-2 Forward | 5'-GCA AAC AGG CTC CGA AGA TAC T-3' |
| Mouse Claudin-2 Reverse | 5'-GAG ATG ATG CCC AAG TAC AGA G-3' |
| Mouse PCNA Forward | 5'-TTT GAG GCA CGC CTG ATC C-3' |
| Mouse PCNA Reverse | 5'-GGA GAC GTG AGA CGA GTC CAT-3' |
| Mouse β-actin Forward | 5'-TGT TAC CAA CTG GGA CGA CA-3' |
| Mouse β-actin Reverse | 5'-CTG GGT CAT CTT TTC ACG GT-3' |

**Table S2 Methods key resources table**

| Items | Company | Source |
|---|---|---|
| *Antibodies* | | |
| β-actin | Sigma-Aldrich | Cat. #: A5316 |
| Alexa Fluor 488 donkey anti-mouse IgG | Invitrogen | Cat. #: A32766 |
| Alexa Fluor 594 donkey anti-rabbit IgG | Invitrogen | Cat. #: A32740 |
| Claudin 2 | Invitrogen | Cat. #: 32-5600 |
| Claudin 5 | Invitrogen | Cat. #: 35-2500 |
| Phospho-NF-κB p65 (Ser276) | Cell Signaling Technology | Cat. #: 3037 |
| Purified Mouse Anti- NF-κB p65 | BD Biosciences | Cat. #: 610869 |
| Phospho-β-Catenin (Ser33/37/Thr41) | Cell Signaling Technology | Cat. #: 9561 |
| β-Catenin | BD Biosciences | Cat. #: 610154 |
| PCNA | Santa Cruz Biotechnology | Cat. #: sc-25280 |
| Villin | Santa Cruz Biotechnology | Cat. #: sc-58897 |
| QTRT1 | Santa Cruz Biotechnology | Cat. #: sc-398918 |
| *Chemicals* | | |
| Fetal bovine serum | GIBCO | Cat. #: 16000044 |
| DMEM | Corning | Cat. #: MT10013CV |
| DPBS | Corning | Cat. #: MT21031CV |
| Penicillin-streptomycin | Corning | Cat. #: MT30002CI |
| TRIzol | Thermo Fisher Scientific | Cat. #: 15596026 |
| iScript cDNA synthesis kit | BioRad | Cat. #: 1708840 |
| SYBR Green PCR kit | BioRad | Cat. #: 1708880 |
| Triton X-100 | Fisher BioReagents | Cat. #: BP151-100 |
| 16% Formaldehyde | Fisher BioReagents | Cat. #: 28908 |
| DAPI | Invitrogen | Cat. #: D21490 |
| Pierce™ ECL Western Blotting Substrate | Thermo Scientific | Cat. #: 32106 |
| Queuine Hydrochloride | Santa Cruz Biotechnology | Cat. #: sc-394021 |
| Dextran sulfate sodium salt | MP Biomedicals | Cat. #: 160110 |
| Vybrant MTT Cell Proliferation Assay Kit | Molecular Probes | Cat. #: V13154 |
| Recombinant Mouse TNF-α | R&D Systems | Cat. #: 410-MT |
| *Software and algorithms* | | |
| FlowJo Version 10.3 | Treestar | www.flowjo.com/ |
| Microsoft Excel Version 16.24 | Office 365 | www.office.com/ |
| Prism Version 8.1.1 | GraphPad | www.graphpad.com/ |
| Zeiss LSM710 confocal microscope | Carl Zeiss | N/A |
| EVOS M5000 Imaging System | Thermo Fisher Scientific | www.thermofisher.com |
| EVOM3 | World Precision Instruments | www.wpi-europe.com |
| Nanodrop 2000 | Thermo Fisher Scientific | N/A |

**Abbreviation list**

CD: Crohn’s disease

DSS: Dextran sulfate sodium

GEO: Gene Expression Omnibus

IBD: Inflammatory bowel disease

IL10: Interleukin 10

PCNA: Proliferating Cell Nuclear Antigen

Q: queuosine

q: queuine

QTRT1: Queuine tRNA-ribosyltransferase 1

QTRT2: Queuine tRNA-ribosyltransferase accessory subunit 2

Q-tRNA: Queuosine-containing tRNA

NF-κB: Nuclear factor kappa-light-chain-enhancer of activated B cells

TEER: Transepithelial electrical resistance

TGT: tRNA-guanine transglycosylase

TJ: tight junctions

TNFα: Tumor Necrosis Factor α

UC: Ulcerative colitis

NOTE: Please save this file locally before filling in the table, DO NOT work on the file within your internet browser as changes will not be saved. Adobe Acrobat Reader (available free here) is recommended for completion.

### The ARRIVE guidelines 2.0: author checklist

#### The ARRIVE Essential 10

These items are the basic minimum to include in a manuscript. Without this information, readers and reviewers cannot assess the reliability of the findings.

| Item | | Recommendation | Section/line number, or reason for not reporting |
|---|---|---|---|
| **Study design** | 1 | For each experiment, provide brief details of study design including: | |
| | | a. The groups being compared, including control groups. If no control group has been used, the rationale should be stated. | Methods: lines 112-202<br>Suppl: lines 4-148 |
| | | b. The experimental unit (e.g. a single animal, litter, or cage of animals). | Methods: lines 112-122<br>Figure legends: lines 544-676 |
| **Sample size** | 2 | a. Specify the exact number of experimental units allocated to each group, and the total number in each experiment. Also indicate the total number of animals used. | Methods: lines 112-122<br>Figure legends: lines 544-676 |
| | | b. Explain how the sample size was decided. Provide details of any *a priori* sample size calculation, if done. | Statistic analysis: lines 205-212 |
| **Inclusion and exclusion criteria** | 3 | a. Describe any criteria used for including and excluding animals (or experimental units) during the experiment, and data points during the analysis. Specify if these criteria were established *a priori*. If no criteria were set, state this explicitly. | Methods: lines 112-122<br>Figure legends: lines 544-676 |
| | | b. For each experimental group, report any animals, experimental units or data points not included in the analysis and explain why. If there were no exclusions, state so. | No exclusions |
| | | c. For each analysis, report the exact value of *n* in each experimental group. | Methods: lines 112-122<br>Figure legends: lines 544-676 |
| **Randomisation** | 4 | a. State whether randomisation was used to allocate experimental units to control and treatment groups. If done, provide the method used to generate the randomisation sequence. | Figure legends: lines 544-676<br>Methods: Statistic analysis: lines 205-212 |
| | | b. Describe the strategy used to minimise potential confounders such as the order of treatments and measurements, or animal/cage location. If confounders were not controlled, state this explicitly. | Methods: lines 112-122 |
| **Blinding** | 5 | Describe who was aware of the group allocation at the different stages of the experiment (during the allocation, the conduct of the experiment, the outcome assessment, and the data analysis). | Authors' contributions: lines 522-526 |
| **Outcome measures** | 6 | a. Clearly define all outcome measures assessed (e.g. cell death, molecular markers, or behavioural changes). | Methods: lines 112-202<br>Suppl: lines 4-148 |
| | | b. For hypothesis-testing studies, specify the primary outcome measure, i.e. the outcome measure that was used to determine the sample size. | Results: lines 216-438 |
| **Statistical methods** | 7 | a. Provide details of the statistical methods used for each analysis, including software used. | Methods: Statistic analysis: lines 205-212 |
| | | b. Describe any methods used to assess whether the data met the assumptions of the statistical approach, and what was done if the assumptions were not met. | Methods: Statistic analysis: lines 205-212 |
| **Experimental animals** | 8 | a. Provide species-appropriate details of the animals used, including species, strain and substrain, sex, age or developmental stage, and, if relevant, weight. | Methods: lines 125-134<br>Suppl: lines 22-40 |
| | | b. Provide further relevant information on the provenance of animals, health/immune status, genetic modification status, genotype, and any previous procedures. | Methods: lines 125-134<br>Suppl: lines 22-40 |
| **Experimental procedures** | 9 | For each experimental group, including controls, describe the procedures in enough detail to allow others to replicate them, including: | |
| | | a. What was done, how it was done and what was used. | Methods: lines 112-202<br>Suppl: lines 4-148 |
| | | b. When and how often. | Methods: lines 112-202<br>Suppl: lines 4-148 |
| | | c. Where (including detail of any acclimatisation periods). | Methods: lines 112-202<br>Suppl: lines 4-148 |
| | | d. Why (provide rationale for procedures). | Methods: lines 112-202<br>Suppl: lines 4-148 |
| **Results** | 10 | For each experiment conducted, including independent replications, report: | |
| | | a. Summary/descriptive statistics for each experimental group, with a measure of variability where applicable (e.g. mean and SD, or median and range). | Results: lines 216-438<br>Figure legends: lines 544-676 |
| | | b. If applicable, the effect size with a confidence interval. | Figure legends: lines 544-676<br>Methods: lines 112-122 |
